# Fast and Robust 2D Inverse Laplace Transformation of Single-Molecule Fluorescence Lifetime Data

**DOI:** 10.1101/2021.01.01.425066

**Authors:** Saurabh Talele, John T. King

## Abstract

Fluorescence spectroscopy at the single-molecule scale has been indispensable for studying conformational dynamics and rare states of biological macromolecules. Single-molecule 2D-fluorescence lifetime correlation spectroscopy (sm-2D-FLCS) is an emerging technique that holds great promise for the study of protein and nucleic acid dynamics as it 1) resolves conformational dynamics using a single chromophore, 2) measures forward and reverse transitions independently, and 3) has a dynamic window ranging from microseconds to seconds. However, the calculation of a 2D fluorescence relaxation spectrum requires an inverse Laplace transition (ILT), which is an ill-conditioned inversion that must be estimated numerically through a regularized minimization. The current methods for performing ILTs of fluorescence relaxation can be computationally inefficient, sensitive to noise corruption, and difficult to implement. Here, we adopt an approach developed for NMR spectroscopy (T1-T2 relaxometry) to perform 1D and 2D-ILTs on single-molecule fluorescence spectroscopy data using singular-valued decomposition and Tikhonov regularization. This approach provides fast, robust, and easy to implement Laplace inversions of single-molecule fluorescence data.

**Significance Statement:** Inverse Laplace transformations are a powerful approach for analyzing relaxation data. The inversion computes a relaxation rate spectrum from experimentally measured temporal relaxation, circumventing the need to choose appropriate fitting functions. They are routinely performed in NMR spectroscopy and are becoming increasing used in single-molecule fluorescence experiments. However, as Laplace inversions are ill-conditioned transformations, they must be estimated from regularization algorithms that are often computationally costly and difficult to implement. In this work, we adopt an algorithm first developed for NMR relaxometry to provide fast, robust, and easy to implement 1D and 2D inverse Laplace transformations on single-molecule fluorescence data.

## Introduction

Single-molecule fluorescence spectroscopy has provided unparalleled access into dynamics of biological macromolecules and the mechanism of biochemical processes (1-3). Typical experimental techniques rely on measuring time-dependent fluctuations in the fluorescence emission wavelength (4-10), intensity (11-13), or lifetime (14-16) from a single chromophore or chromophore pair. The dynamics of biochemical processes of interest are then inferred from the analysis of the photon stream. The most widespread technique is FRET (sm-FRET) (1-3, 17, 18), which leverages the highly sensitive distance dependence for dipole-dipole coupling between two chromophores to monitor nanometer scale motions of a biomolecule. Conformations are distinguished by the emission wavelength, which is either from the acceptor or donor chromophore depending on the separation distance. Transitions between two conformations are observed as abrupt changes in emission intensity at both the acceptor and donor emission wavelength. However, the need for site-specific labelling of two chromophores on a single biomolecule makes its application to proteins challenging (17), in particular when compared to nucleic acids (18), which are relatively easy to modify. Furthermore, the temporal resolution is typically limited to 10-100 ms (19) as the low photon flux from a single-molecule requires temporal binning in order to obtain a meaningful analysis of the photon stream.

Single-molecule 2D fluorescence lifetime correlation spectroscopy (sm-2D-FLCS), first developed by Tahara and coworkers (20-22) and later extended into the single-molecule regime by Schlau-Cohen and workers (23), provides an alternative analysis of single-molecule data that does not sacrifice chemical selectivity or temporal resolution. In this approach, time-correlated single photon counting (TCSPC) is used to detect fluorescence intensity and emission delay time from a single chromophore, either freely diffusing in dilute solution (20-22) or surface immobilized (23). The data are recorded in the TTTR mode which generates a real-time photon stream (characterized by a global macro time) with a recorded emission delay time (characterized by a micro-time) for each registered photon (24). Distinct chemical species in the system are distinguished by their fluorescence lifetime. Thus, chemical exchange between two states can be kinetically resolved on the condition that they have different fluorescence lifetimes.

A series of 2D photon histograms are generated by cataloging all photon pairs separated by a systematically varying waiting time *ΔT* from an experimentally recorded photon stream (20, 21). A 2D inverse Laplace transform (2D-ILT) of the lifetime histogram generates a 2D-FCLS spectrum at the given *ΔT*. Analogous to 2D-NMR (25), species that do not undergo any form of exchange during *ΔT* appear as diagonal peaks in the 2D spectrum, while species that exchange during *ΔT* appear as cross-peaks. Measuring 2D spectra for a series of *ΔT* allows the overall correlation function to be effectively split into its components comprising auto-correlations for the diagonal components and cross-correlations for the off-diagonal components. Thus, the chemical exchange kinetics among the components can be measured directly through time correlation functions reflecting chemical exchange between two states (20-23). This is an important advancement in single-molecule spectroscopy as 1) the technique requires only a single chromophore whose lifetime is sensitive to the environmental surroundings, 2) resolving the lifetime spectrum over two-dimensions allows the forward and reverse transitions to be measured separately, as they are recorded in different regions of the spectrum, and 3) the dynamic range extends from microseconds to seconds. Furthermore, as the technique requires no additional optical setup or heavy computing hardware, it can be implemented on any setup already used for FCS or FRET.

The challenge of this experimental approach is the need to perform a 2D-ILT, which is an ill-conditioned problem and is therefore numerically unstable. This results in multiple solutions satisfying the same problem when solved with traditional least-square analysis (26). Instead, this class of problems must be solved via regularized least-square analysis, which imposes a penalty on solutions with undesired features (27). To date, the calculation of the 2D spectra is traditionally performed using the maximum entropy method (MEM) (20-23), which produces a solution obeying Bayesian statistics. The application of MEM has two severe limitations: First, it involves a constrained optimization since entropy cannot have a negative value, so one must check for the positivity constraint for each iteration and modify the fit accordingly. This makes the approach computationally inefficient. Second, the 2D spectra is fit as a lexicographically ordered 1D vector resulting in a fitting kernel matrix whose size is of the order of the 4^th^ power of number of points in the lifetime spectra, leading to a tremendous computational cost. To decrease the computational cost the fitting kernels are often logarithmically binned, which results in decreased spectral resolution (21).

The challenge of performing 2D inverse Laplace transforms (i.e. converting the time dependent relaxation data to a relaxation rate spectrum) is not unique to fluorescence spectroscopy. Indeed, multi-dimensional Laplace inversions are commonly employed in NMR spectroscopy, for instance in the computation of T1-T2 correlation spectrum from relaxometry data. Work by L. Venkataramanan and coworkers (28) outlined an efficient algorithm for Laplace inversion in NMR relaxometry data that leveraged singular-valued decomposition (SVD) and Tikhonov regularization. Subsequent improvements of this approach have been reported (29, 30). In this paper, we adopt this general approach for application to single-molecule fluorescence spectroscopy analysis. This algorithm reduces the computation time to merely a few seconds per 2D spectrum, and hence enables the analysis of large data sets with high resolution.

## Methods

A 2D-FLCS spectrum is generated from a 2D-ILT of a lifetime correlation histogram, easily measured using standard time-correlated single-photon counting (TCSPC) techniques (20, 21). We start with a familiar 1D picture to introduce the concept of ILT. Given a time series of recorded photons, the emission delays are distributed exponentially and can be represented as

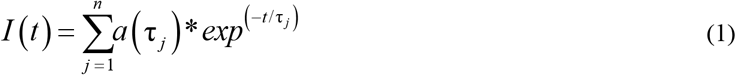

for *n* independent components with amplitudes *a*_*j*_ and respective fluorescence lifetimes *τ*_*j*_. This can be generalized for the sake of broader application as

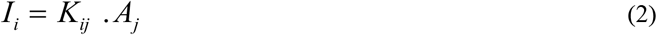

Where *K* is a kernel with pre-defined basis of fluorescence lifetimes *K*_*ij*_ *= exp(-t*_*i*_*/τ)*, and *A*_*j*_ is a column vector representing the amplitudes corresponding the species *τ*_*j*_. Kernel *K* can be defined such that the experimental lifetimes are included in [*t*_*min*_, *t*_*max*_] with sufficient resolution such that any measured decay can be represented by a pre-calculated kernel and an amplitude vector. Under this construction, *A(τ*_*j*_) represents a 1D ILT of *I(t)* as we convert the time dependent decay into its lifetime components and their probability distribution amplitudes.

This can be easily extended to 2D. Cataloging all the photon pairs separated by a lag time *ΔT* in the experimentally recorded time series and having emission delays *t*_*1*_ and *t*_*2*_, we can generate a 2D histogram with axes as *t*_*1*_ and *t*_*2*_, where each point *M(t*_*1*_, *t*_*2*_, *ΔT)* represents the number of coincidences when a photon with delay *t*_*2*_ is detected *ΔT* after having detected a photon with delay *t*_*1*_.

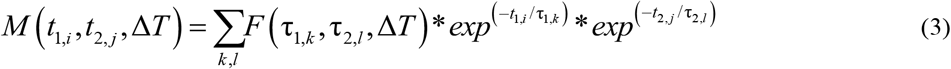

Using the kernel structure mentioned above, we can write in matrix notation

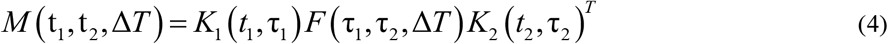

where *F(t*_*1*_, *t*_*2*,_ *ΔT)* is the joint probability distribution of occurrence of a photon with lifetime *τ*_*2*_ occurring *ΔT* after registering a photon with lifetime *τ*_*1*_. Each point on *F(t*_*1*_, *t*_*2*,_ *ΔT)* represents the amplitude of correlation between the components at (*τ*_*1*_, *τ*_*2*_) separated by lag time *ΔT*, the auto-correlations appear along the diagonal of *F(t*_*1*_ *= t*_*2*_) and the cross correlations appear as off-diagonal peaks of *F(t*_*1*_ ≠ *t*_*2*_). By varying *ΔT*, one can determine the separated time correlation functions of the constituent components in any fluorescence emission time series.

Determining *F(ΔT)* is equivalent to performing a 2D-ILT on *M*. For efficient handling, we can convert the 2D form of Eq.(4) to 1D by lexicographically ordering the matrices *m = vec[M], f = vec[F]* and can write the kernel operations *K*_*1*_ and *K*_*2*_ by a single operator *K*_*0*_ given by the Kronecker product.

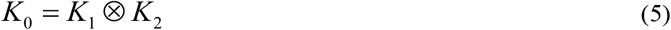

And we represent the equivalent 1D problem as:

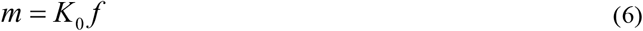

Now, the only task at hand is to solve for *f* given *m* and *K*_*0*_. As previously mentioned, this equation cannot be solved analytically since it is an ill-conditioned inversion resulting in a solution that is not stable or unique. Instead, regularized least-squared technique must be used to approximate the inversion. In regularized least square minimization, the least square minimizes the difference between the input data and the fit, while the regularization imposes a penalty on undesired features of the fitted solution. The regularization prevents overfitting or numerical instability. To avoid any bias, we start with a uniform guess of f and iteratively find the solution by minimizing the objective function given by

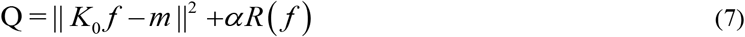

where ║.║^2^ is the Frobenius norm and represents the least square term, *R(f)* is the regularization function of choice, and *α* is a regularization constant that weighs the importance of the least square fitting vs regularization. The choice of *α* is critical to appropriate fitting; too small and the minimization remains unstable, too large and fit may not reflect the underlying experimental data. Two regularization methods discussed here include the commonly employed maximum entropy method (MEM) (31-33) and Tiknokov regularization (34, 35). Ultimately, we highlight the strength of Tikhonov regularization at providing a fast and robust method for performing 2D-ILT.

### 2D-ILT by Maximum Entropy Method

The maximum entropy method (MEM) is commonly employed fitting technique used for a number of applications, including image reconstruction (31, 32) and spectroscopy (33). It was also the approach applied by Tahara and coworkers in their original development of 2D-FLCS spectroscopy (20, 21). MEM uses a regularization function based on Shannon entropy penalization *ϕ(s) = -s*log*s*. The regularization function can be written as,

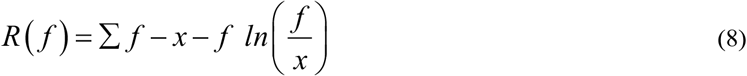

where *x* represents the estimated prior fit of the experimental data. The algorithm is initiated by taking *f* to be a flat distribution and then optimized according to Eq.(7). Fitting is typically initiated with a large *α* parameter, which is iteratively decreased until the classical MEM condition is satisfied (36). This method ensures that off all the possible solutions, the solution with the maximum information entropy is chosen.

For typical 2D-FLCS spectra the data sets are too large to compute ILT using MEM efficiently. Instead, the data is binned logarithmically, which decreases the data size at the cost of spectral resolution. Furthermore, to obtain a meaningful ILT using MEM, one must impose a non-negativity constraint after every iteration, since the entropy cannot be defined for negative values. This significantly decreases the efficiency of the calculation.

### 2D-ILT by Tikhonov Regularization

An alternative approach has been developed in NMR spectroscopy based on SVD based data compression and Tikhonov regularization (28-30). This approach differs from MEM in several ways. First, the size of the numerical calculation is greatly reduced by compressing the data using singular-valued decomposition (SVD) instead of non-uniform binning. Since the kernels are smooth functions, the elementwise data is vastly redundant and SVD can reduce the data size by roughly a thousand-fold without compromising the quality of the fit or the spectral resolution. Second, it uses Tikhonov regularization on the compressed data, which ensures a unique solution to the optimization due to the quadratic nature of the terms. Third, it employs Butler-Reeds-Dawson (BRD) method to transform the constrained optimization to an unconstrained optimization which is computationally efficient to implement. Lastly, it provides a method for choosing an appropriate regularization constant scaled with the noise variance of the data. The outline of the method follows what is presented in references (28, 30).

### Data compression

To compress the size of the kernel functions we perform a SVD using the built-in MATLAB subroutine *svds(K, n)*, where *K* is the kernel matrix and *n* is the number of singular values chosen and is represented in matrix form as:

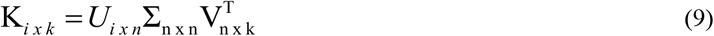

Here, *U* and *V* are unitary matrices, and ∑ is a diagonal matrix with the singular values in descending order along the diagonal. Our data matrix from Eq.(4) can be compressed as

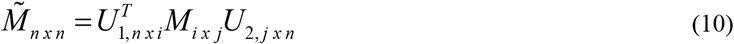

and kernels *K*_*1*_ and *K*_*2*_ can be compressed as:

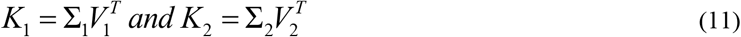

Converting this to 1D by lexicographic ordering, we get 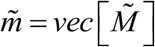 and *K*_0_ = *K*_1_ ⊗ *K*_2_ This compression allows us to solve the inversion with a significantly smaller data set:

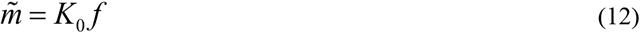

### Tikhonov regularization

Using Tikhonov regularization, the objective function to be minimized from equation (7) becomes

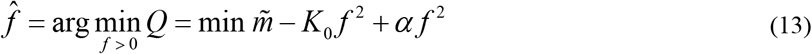

The quadratic nature of the terms ensures a unique solution. The second term penalizes solutions that have large norms, characteristic of functions with sharp features. Solutions with smoothly varying features are therefore promoted, and *α* scales the desired smoothness with respect to the least squares difference. However, since *f* is the probability distribution, it cannot take negative values which makes this a constrained optimization problem. The convert the inversion to an unconstrained problem the Butler-Reeds-Dawson (BRD) algorithm is employed (28). We invoke a vector *c* which maps *f* as

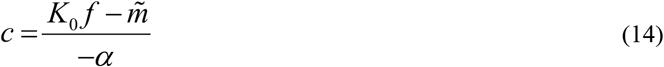

The unconstrained problem then becomes

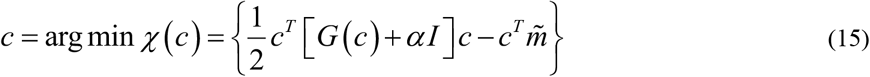

where,

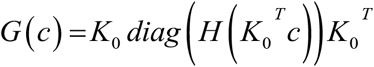

Here, *H(·)* denotes the Heavyside function, which ensures positive semi-definiteness. The minimization of *χ(c)* can be carried out via standard unconstrained inverse Newton minimization routines such as *fminunc* in MATLAB. The required gradients and Hessian are easily computed as

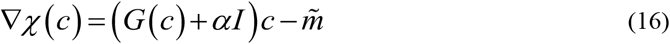

and,

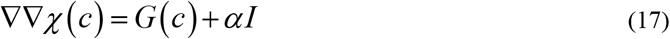

Using the optimized vector *c*, the ordered vector *f* is calculated as

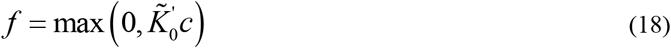

The 2D-ILT spectrum *F* is then given by reshaping the *f*_*1xkl*_ vector to a matrix *F*_*kxl*_. For choosing the *α* parameter, we start with a large value of *α* to estimate *c*. The recommended optimum *α* is given by

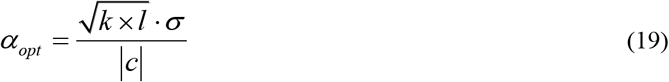

with,

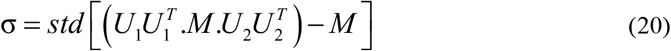

The next vector *c* is iteratively evaluated using this *α*_*opt*_. A suitable convergence criterion is the relative difference between consecutive *α* to be less than 0.1%.

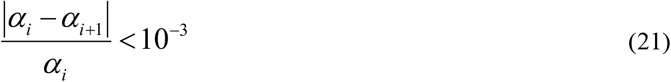

Typically, convergence is achieved in around 10 iterations.

## Results and Discussion

We demonstrate the application of this method using Markovian Monte carlo simulations to generate artificial photon time series for a two-state system with user defined inputs for fluorescence lifetime, emission intensity and transition rate matrix (**see Supplemental Information for details**). We compare the compression efficiency, tolerance to noise, and the timing of the algorithm for various relevant parameters.

### Implementation

We simulate a two-state system with fluorescence lifetimes of *τ*_*1*_ = 1.0 ns and *τ*_*2*_ = 3.0 ns, and exchange rates of *k*_*f*_ = *k*_*r*_ = 1×10^3^ s^-1^ corresponding to an exchange time of 1 ms. The corresponding 1D and 2D lifetime spectra were obtained by ILT by Tikhonov regularization as described above, using a basis set of size *L* = 100 and with a singular valued decomposition using 20 greatest values. The 1D photon histogram, resulting fit computed by a 1D-ILT, and residual are shown in **Fig. 1B, C**. The 1D relaxation rate spectrum shows two peaks located at *τ*_*1*_ = 1.0 ns and *τ*_*2*_ = 3.0 ns, consistent with the input parameters of the simulation (**Fig. 1D**).

**Figure 1.**
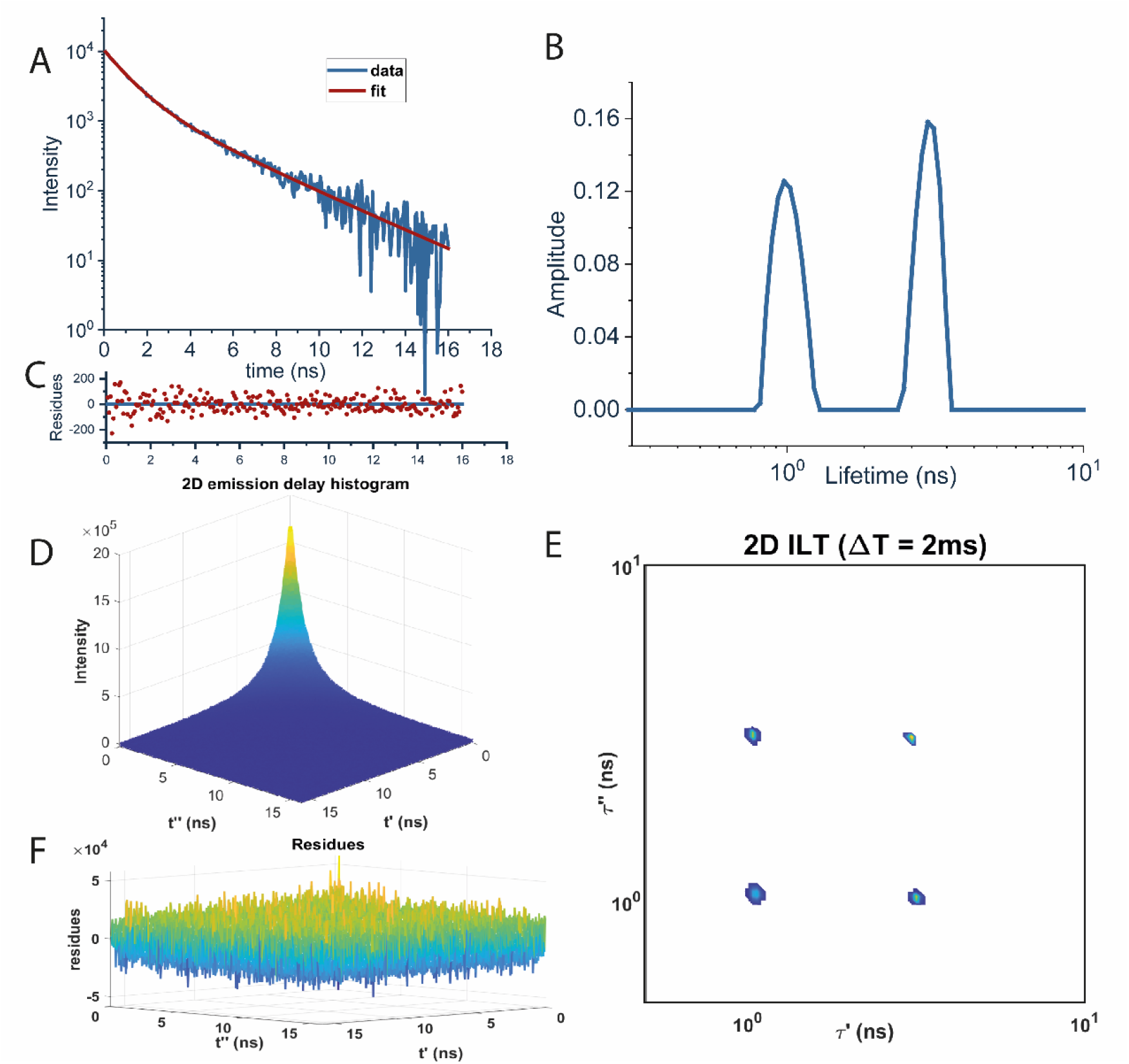
Laplace inversions by Tikhonov regularization. Monte carlo simulations generate an artificial photon stream of a two component system with fluorescence lifetimes of *τ*_*1*_ = 1 ns and *τ*_*2*_ = 3 ns undergoing equilibrium chemical exchange at a rate of 1×10^3^ s^-1^. (**A**) Raw 1D-photon histogram (blue) and the obtained fit (red) from 1D-ILT. (**B**) The 1D lifetime spectrum shows two peaks with relaxation rates of 1 ns and 3 ns, consistent with fluorescence lifetime of the two components in the system. (**C**) Residual between the 1D photon histogram and the fit obtained by 1D-ILT. (**D**) 2D photon correlation histogram and (**E**) 2D relaxation rate spectrum computed at a waiting time of *ΔT* = 2 ms obtained from a 2D-ILT. The spectrum consists of diagonal peaks at 1 ns and 3 ns, as well as cross-peaks indicating chemical exchange between the two species. (**F**) Residual between the 2D photon correlation histogram and 2D-ILT fit.

The 2D photon correlation histogram evaluated at *ΔT* = 2 ms, resulting fit computed by a 2D-ILT, and residual are shown in **Fig. 1E, F**. A 2D-FLCS spectrum shows two diagonal peaks at *τ*_*1*_ = 1.0 ns and *τ*_*2*_ = 3.0 ns, and two off-diagonal cross-peaks between these transitions (**Fig. 1G**). The cross-peaks indicate that chemical exchange has occurred within the timescale *ΔT* = 2 ms, which is consistent with the input transition rate matrix.

Computing the 2D-FLCS spectrum for a series of *ΔT* delay times allows the chemical exchange kinetics to be directly measured. **Fig. 2A-C** shows the 2D-FLCS spectrum measured for delay times ranging from 10 μs to 2 s. At early times (*ΔT* < *τ*_*exchange*_), only diagonal peaks are observed as no chemical exchange had occurred (**Fig**.**2A**). As *ΔT* is increased, cross-peaks emerge as chemical exchange occurs (**Fig**.**2B**). The exchange cross-peaks reach a steady state as *ΔT* > *τ*_*exchange*_ (**Fig**.**2C**). Monitoring the amplitude of the diagonal and cross-peaks provides direct information regarding the chemical exchange process. The amplitude of the peaks can be expressed as,

**Figure 2.**
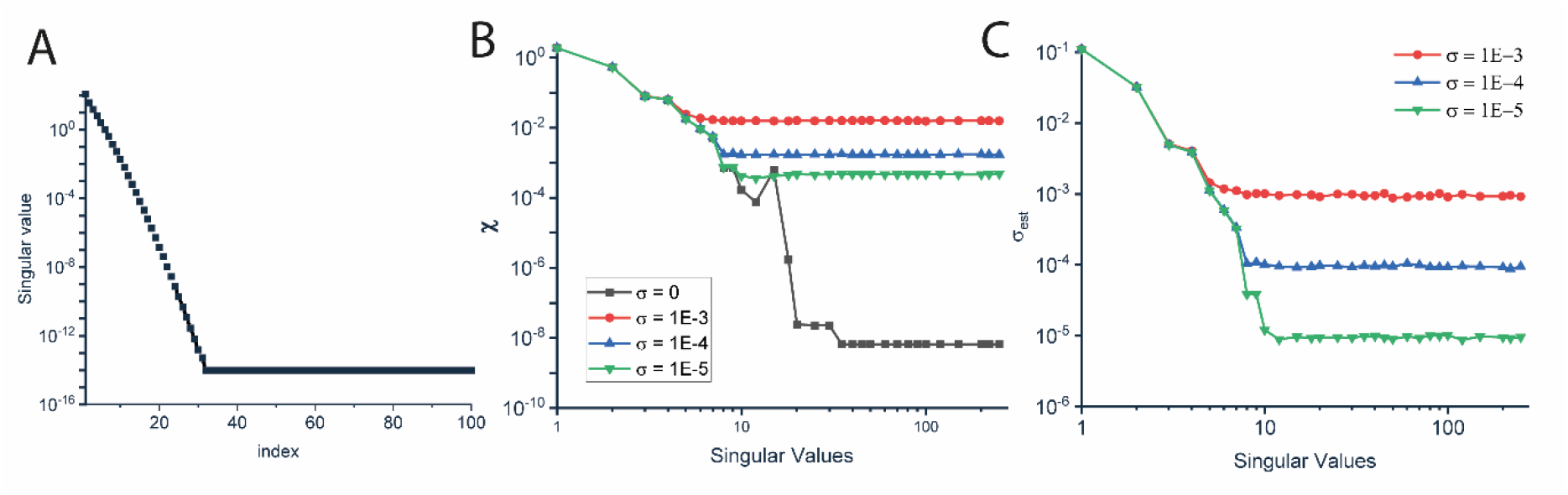
Chemical Exchange kinetics of a two-state system. 2D-FLCS spectra computed at (**A**) *ΔT* = 10-20 μs, (**B**) *ΔT* = 100-200 μs, (**C**) *ΔT* = 1-2 ms. At *ΔT* < *τ*_*exchange*_, no cross-peaks are observed between the two states, indicating no exchange has occurred. As *ΔT* approaches *τ*_*exchange*_ cross-peaks emerge indicating both forward and reverse transitions between the two states. (**D**) The kinetics of the exchange process are reflected in the auto-correlation and cross-correlation functions of the diagonal and off-diagonal peaks, respectively. Fitting either the auto-correlation or cross-correlation functions provide a direct measure of the exchange kinetics in the system.

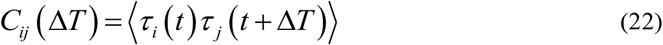

Where *i* and *j* represent the state of the system. The diagonal is given by the condition *i = j*, while cross-peaks are given by the condition *i* ≠ *j*. The two auto-correlation and cross-correlation functions obtained from the 2D-FLCS spectra are shown in **Figure 2D**. As expected, the auto-correlation functions decay on the timescale of *τ*_*exchange*_, while the cross-correlation functions grow on the timescale *τ*_*exchange*_. Fitting the data provides a direct measure of *τ*_*exchang*_ for the system.

In practice, lifetime histograms are convoluted by a systematic instrument response function IRF and do not manifest as pure exponential decays. We can account for this by fitting the data with kernels convoluted by a known IRF which can be measured experimentally from scattering of the excitation pulse from a colloidal medium. The experimentally observed TCSPC histogram is modelled from Eq.(1) as

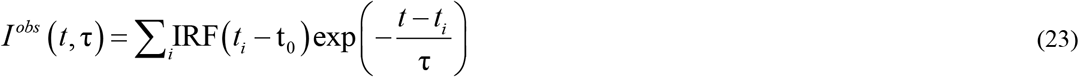

where, *t*_*0*_ is the unavoidable zero-time shift for the IRF. It is important so estimate the correct value of *t*_*0*_ as it can lead to undesirable errors in the ILT spectrum. To circumvent this, we calculate the 1D-ILT spectrum at various values of zero-time shifts and use the range over which the fitting error *χ* for the is minimum. The ILT obtained from this range is then averaged to obtain the final 1D-ILT spectrum to be used for further analysis. 2D spectrum is calculated using the same range of zero-time-shifts. The kernels to be fitted are also convoluted with the IRF similarly.

The fitting was performed on an Intel-Core i7-4790 CPU and using the code written in MATLAB 2020a software. The table below shows the CPU time required for execution of the fits for various sizes of basis set.

| Basis Size | 1D ILT time (seconds) | 2D ILT time (seconds) |
| --- | --- | --- |
| 1000 | 0.4 | 1000 |
| 100 | 0.1 | 7 |
| 50 | 0.05 | 1 |

### Compression efficiency

Data compression increases computational efficiency by reducing the size of the problem. However, there is a tradeoff between the amount of compression and loss of information. The measure of compression in our case is given by the number of singular values chosen to represent the kernel of exponentials as well as the data. **Figure 3A** shows the singular values for a kernel comprising of 1000 basis exponentials in descending order. It is observed that only the 30 greatest singular values are enough to sufficiently describe the kernel. The required number of singular values decreases with basis size. The effect of compression can be quantified by calculating the estimated noise variance **Figure 3B** and the norm of residuals after the fitting **Figure 3C**. We notice that both the estimated noise variance and norm of residuals plateau at certain levels proportional to the input noise magnitude (*SV* = 5, *σ* = 1×10^−3^; *SV* = 8, *σ* = 1×10^−4^; and *SV* = 12, *σ* = 1×10^−5^) indicating that the quality of fit does not significantly improve on using more values than *SV*_*plateau*_. However, fitting with less values than *SV*_*plateau*_ can lead to the amplification of noise and artifacts in the ILT spectrum. We choose SV = 20 as a general parameter in the analysis which we find optimal considering the computational time and minimal loss of information since the *χ* and *σ*_*est*_ both plateau before that value even without any input background noise (the limit is dictated by numerical precision).

**Figure 3.**
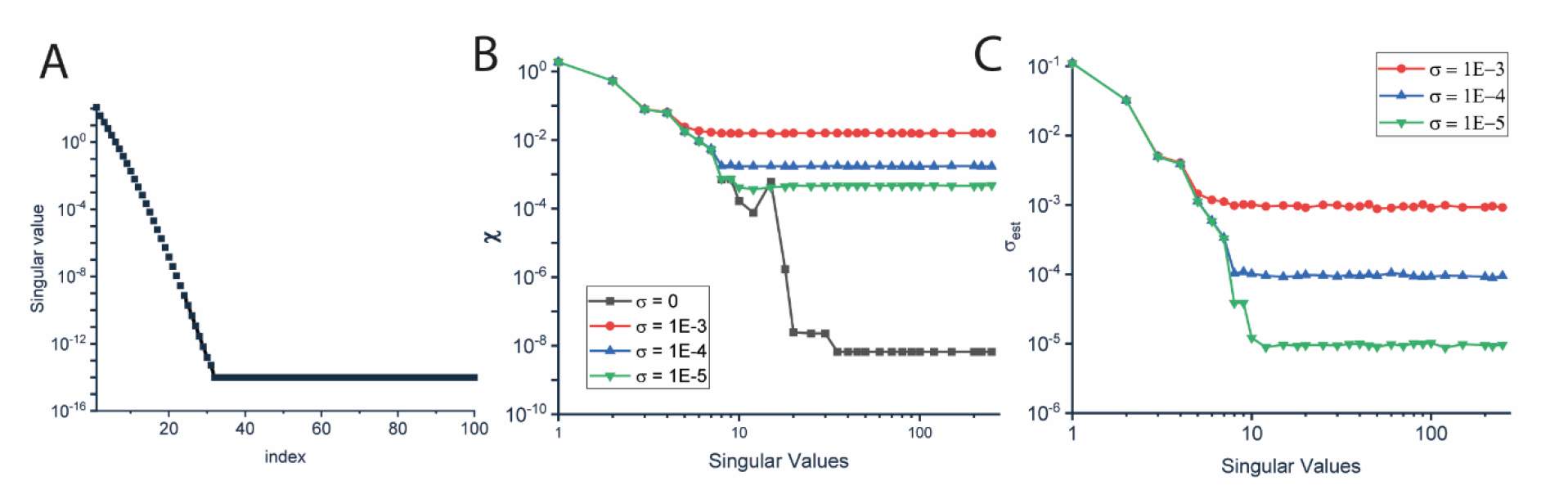
Characterization of the effect of SVD on ILT. (**A**) Singular values of the kernel matrix comprising of exponential decays with 100 distinct lifetimes in range (0.1 ns, 10 ns). (**B**) Residual of fit (χ) plotted against the number of singular values used at various levels of background noise (*σ*). (**C**) Estimated background noise in the data calculated before fitting, predicted from (1.21). Using fewer than 10 singular values leads to amplification of the background noise. However, all the information about the input data is captured under 20 singular values.

### Effect of noise

The presence of experimental noise can affect the spectral resolution and the width of detected peaks. The two major sources of noise in fluorescence experiments are the background noise (normally distributed) and noise due to finite photons in the TCSPC histogram. The effect of noise in the fitting directly manifests as a smoothening bias to the *α* parameter, since the variance of the noise is included in the convergence criterion (20). Larger noise variance thus causes *α* to converge at higher values to prevent overfitting and one observes broadening of the fitted peaks. This is demonstrated in **Figure 4A**, which shows the log-log plot of *χ* vs *α* (also known as the S-curve for its shape) for various levels of SNR. It is recommended to choose the *α* at the “heel” of this S-curve, which is observed to be close to the regularization value predicted by the algorithm. The *α*_*opt*_ suggested by the algorithm for those levels is shown by the red dot. **Figure 4B** shows *α*_*opt*_ as a function of background noise. **Figure 4C, E** shows the 1D and 2D ILT fits for recommended *α*_*opt*_ corresponding to the background noise level (*σ* = 3×10^−3^). **Figure 4C** also shows 1D ILT overfit (orange spectrum) and underfit (green spectrum), constructed by forcing the alpha to value *α*_*overfit*_ = 10^−7^ and *α*_*underfit*_ = 10^−1^. This highlights the importance of correctly choosing *α*. Similarly, **Figure 4D, F** show the 2D ILT underfit and overfit for the fixed variance of background noise.

**Figure 4.**
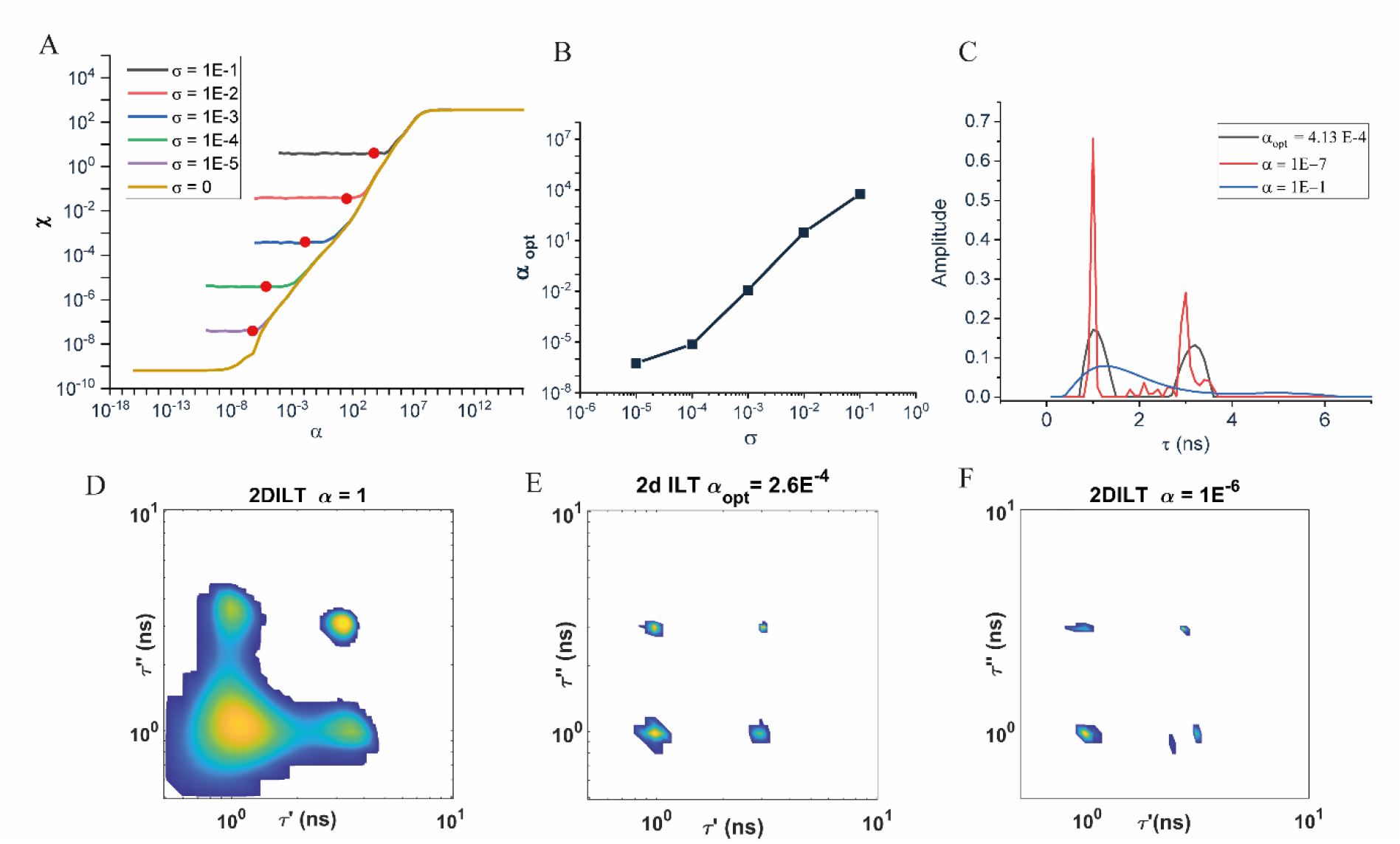
Characterization of the effect of *α* on ILT. (**A**) A log-log plot of *χ* vs *α* for various levels of background noise shows an characteristic S-curve. The red dots shown indicate the optimum *α* suggested by the algorithm. (**B**) *α*_*opt*_ plotted as a function of *σ*. (**C**) Three representative 1D-ILTs of a two state system with lifetimes τ_1_ = 1 ns, τ_2_ = 3 ns and background noise σ = 1×10^−3^ computed with different *α* values. The strong dependence of the obtained spectrum on the *α* demonstrate the importance of proper selection of *α*_*opt*_. At *α* = *α*_*opt*_, we observe two peaks in the 1D-ILT (black) at the corresponding lifetimes of 1 ns and 3 ns. For *α* > *α*_*opt*_ (blue), we observe an underfit spectrum with no resolved features. For *α* < *α*_*opt*_ (red), we observe two prominent peaks at 1 ns and 3 ns along with false peaks due to overfitting. Similarly, (**D, E, F**) show the effect of choice of *α* for 2D-ILT with (**D**) underfitting (*α* > *α*_*opt*_), (**E**) optimal fitting (*α = α*_*opt*_), and (**F**) overfitting (*α* < *α*_*opt*_).

## Conclusion

Single-molecule 2D-FLCS is a powerful tool for studying dynamics of biological macromolecules, though the difficulty of computing inverse Laplace transforms may be prohibitive to its wide-spread application. Here, we outline a fast and robust method of computing 2D inverse Laplace transforms. The method, based on singular-valued decomposition and Tikhonov regularization, is adopted from NMR spectroscopy for application to single-molecule fluorescence spectroscopy. The approach allows for stable inversions of large, noisy data sets, common in single-molecule spectroscopy, to be carried out efficiently, without sacrificing the resolution of the spectra. Furthermore, using Monte carlo simulations to generate artificial photon streams, we demonstrate that this technique is robust in terms of the spectral resolution, noise tolerance, and computational efficiency. This provides an alternative method (beyond typical maximum entropy methods) for performing Laplace inversions of single-molecule fluorescence data that is easily implemented.

