## Supporting Information for "Fast and Robust 2D Inverse Laplace Transformation of Single-Molecule Fluorescence Lifetime Data"

**Methods**

1. **Workflow**
2. **Monte carlo simulations**
3. **Instrument response function**

**Figures**

### 1. Experimental Workflow

Here we describe stepwise procedure of data analysis for the sm-2D-FLCS application

i. **PTU_read:** From TCSPC experiment, select the single molecule trajectories. for PicoQuant TCSPC setup, the files are stored with '.ptu' extension. We use a demo code 'Read_PTU.m' to convert the files to .csv to be used in further analysis [1]. The data stored is in the form of photon index, its global arrival time (MacroTime) and its emission delay time after excitation (MicroTime).

ii. **Partitioning:** for 2D FLCS, we need to evaluate the 2D emission delay histogram at various lag times ( $\Delta T$ )[2]. We employ the following procedure

a. For a given $\Delta T$, define a window of size $\Delta\Delta T$ such that its upper bound is $\Delta T + \frac{\Delta\Delta T}{2}$ and the lower bound is $\Delta T - \frac{\Delta\Delta T}{2}$.

b. Initialize a 2D empty array (M) of size (N x N), where N is the number of TCSPC channels.

c. For the first photon in the photon trace, store its macrotime (as x).

d. Find all the photons in the trace whose macrotime lies within the constructed window.

$$\Delta T - \frac{\Delta\Delta T}{2} < T_M < \Delta T + \frac{\Delta\Delta T}{2} \quad (1)$$

e. Store all the recorded macrotimes as an array.

f. In the 2D array, in the column denoted by x, increment the elements by 1 for each occurrence of their corresponding macrotime in array.

g. Repeat steps [c] to [f] for all the photons in the trace.

h. Repeat steps [a] to [g] for all $\Delta T$ windows.

iii. **ILT analysis: [3]**

a. Define a basis with an upper bound lower bound and number of points.

b. Import data and IRF

c. Generate a kernel matrix (whose size is length of histogram x ILT basis size).

$$K_{ij} = exp^{\left(-t_i/\tau_j\right)} \tag{2}$$

Convolute the kernel with IRF.

d. Compute the SVD of the kernel and stored the compressed U, S, V matrices

the compressed kernels are $K_1 = \Sigma_1 V_1^T \; and \; K_2 = \Sigma_2 V_2^T$.

Generate the Kronecker product matrix $\tilde{K}_0 = \tilde{K}_1 \otimes \tilde{K}_2$.

e. Unconstrained optimization: initialize a constant array (c), start with initial α

$$\chi(c) = \frac{1}{2} c^T \left[G(c) + \alpha I\right] c - c^T \tilde{m} \tag{3}$$

$$\nabla\chi(c) = \left(G(c) + \alpha I\right) c - \tilde{m} \tag{4}$$

$$\nabla\nabla\chi(c) = G(c) + \alpha I \tag{5}$$

Where $G(c) = K_0 \, diag\left(H\left({K_0}^T c\right)\right) {K_0}^T$ and H is the Heaviside function.

f. $\alpha_{opt} = \frac{l.\sigma}{|c|}$. where l is the size of the basis, σ is noise variance.

g. Check convergence condition: $\left|\frac{\alpha_i - \alpha_{i+1}}{\alpha_i}\right| < 10^{-3}$ (6)

h. $F_{ILT} = max(0, K^T c)$

### 2. Monte Carlo Simulations

<u>Input Parameters</u>

| Parameter | Value | Comment |
|---|---|---|
| Simulation time step | 25 ns | 1/Laser rep. rate (40MHz) |
| No. of TCSPC channels | 256 | Width = 64 picoseconds |
| Number of conformations | 2 | |
| Brightness | $[10^4, 10^4]$ | photons/second |
| Fluoresce lifetimes | [1, 3] | nanoseconds |
| Transition probability matrix | | |
| Simulation size | 50 seconds | $2\,x10^9\ time\,steps$ |

The monte carlo simulation is a 4-step process: Initialize, Reaction, Photon emission and Detection.

Reaction: The chemical reaction is governed by transition rate matrix. The initial states are randomly chosen based on the steady state probabilities. In case of a 2 state simulation, $P_1 = K_{2\to1} / \left(K_{1\to2} + K_{2\to1}\right)$ and $P_2 = K_{1\to2} / \left(K_{1\to2} + K_{2\to1}\right)$. The initial state is stored, and the probability of transition is calculated as $P_{i\to j} = K_{i\to j}.\delta t$ where $K_{i\to j}$ is the rate of transition from state $i$ to $j$ and $\delta t$ is the time step. A uniform random number $R_r$ is generated between (0,1) ('*rand()*' function in MATLAB) and compared with $P_{i\to j}$. The reaction succeeds if $R_r < P_{i\to j}$ and the state is updated from $i$ to $j$. Otherwise the molecule remains in $i$ state. The updated state is stored, and the process is repeated for the length of simulation using the new initial state as the updated state. The result is an array which shows the trajectory of chemical reaction of the molecule.

Photon emission: Each state is assigned a brightness property $B_i$ in units of $photons / second$ and a characteristic fluorescence lifetime $\tau_i$ .For every point along the reaction trajectory generated above, the probability of emission is given by the $P_{em} = B_i.\delta t$ . $P_{em}$ is compared with another uniform random number $R_{em}$ between (0,1) and photon emission is registered during the time step if $R_{em} < P_{em}$. This results in a photon emission trajectory. For every emitted photon, an emission delay time is generated from using a exponential random number generator (*'exprnd(mu)'* in MATLAB) with the lifetime equal to the lifetime of current state. The result is an emission delay trajectory corresponding to the photon emission trajectory.

Detection: The photon emission trajectory is called the MacroTime $(T)$ and the emission delay trajectory is the MicroTime $(t)$. The microtime is discretized based on the expected TCSPC parameters (channel width = 64 picoseconds and 256 bins. Photon data beyond the bin limit (16 ns = 256 x 64ps) is discarded. The final resultant photon stream is stored in the form of an array with 2 columns – MacroTime and MicroTime $(T_i, t_i)$

### 3. Instrument response function (IRF)

Experimentally, the emission delays are influenced by the instrument response of the detectors and the electronics involved in the detection. The instrument response for TCSPC can be measured by recording an emission delay histogram from a scattering sample for example LUDOX silica beads. Deconvolving the data to separate the delays from fluorescent sample and instrument response is numerically difficult process. A simple alternative is to convolve the fitting kernels with the instrument response and fit it to the experimentally convoluted data.

For simulations, we model the instrument response as

$$IRF(t,t_0) = \frac{\sqrt{\pi} w A}{2} exp\left(\frac{t_0 - t}{\tau_k} + \frac{w^2}{4\tau_k^2}\right) Erfc\left(\frac{t_0 - t}{w} + \frac{w}{2\tau_k}\right) \quad (7)$$

Where $w$ is the width,

$A$ is the amplitude factor, $t_0$ is the zero-time shift and $\tau_k$ are the lifetimes.

We generate a random number distribution using rejection sampling to be added to the emission delays.

Ideally, one expects to have constant zero-time shifts for the data and IRF. However inaccurate measurement can cause significant errors in the positions of the peaks in the estimated lifetime spectra. To circumvent this problem, we adjust the zero-time shifts in a narrow range around the expected value and monitor the $\chi^2$ error w.r.t the shift. The value of zero-time shifts where the error is minimum is chosen to be used as the final zero-time shift for the IRF for the analysis. Figure1 shows the simulated emission delay histogram after IRF convolution and the zero-time shift error.

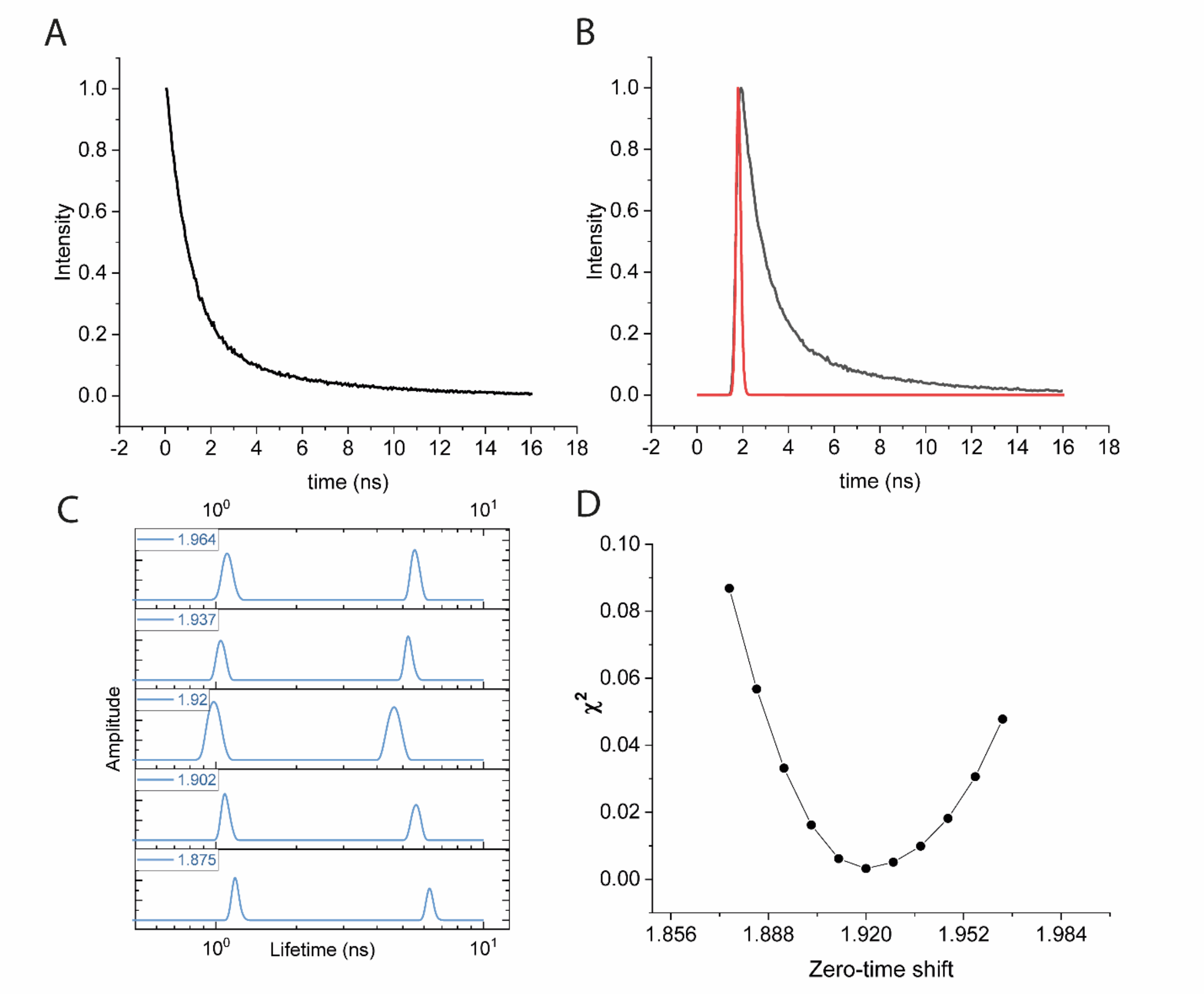


**Figure 1** Monte carlo simulation: A shows the 1D emission delay histogram. B shows the emission delay histogram (black) convoluted by the Instrument response function (red). C shows the 1D ILT spectra at various zero-time shifts of the IRF. D shows the evaluated least-square error in the fit and convoluted histogram. The ILT spectra at the minima of the plot in D is selected as the 1D lifetime spectra.
